# Age-dependent changes in adult fat content differ between the vinegar fly, *Drosophila melanogaster,* and its parasitoid amber wasp, *Leptopilina heterotoma*

**DOI:** 10.64898/2026.06.24.733671

**Authors:** Mathilde Scheifler, Maude Quicray, Caroline M. Nieberding, Bertanne Visser

## Abstract

Fat reserves are critical for sustaining life and insects generally accumulate fat when sufficient sugars and other carbohydrates are consumed. Most studies on insect fat reserves measure fat content at a single age, early in adult life, while fat content is expected to change as insects age. In this paper, we compared age-dependent changes in fat percentages during adulthood (0-56 days after emergence) in lean and fat *Drosophila melanogaster* females and in females of 5 inbred lines of the parasitoid wasp *Leptopilina heterotoma,* reared on lean and fat hosts. We expected *D. melanogaster* to accumulate fat irrespective of teneral fat content at emergence. Fat *D. melanogaster* females, however, lost fat during the first week of life, while lean flies did not. Both lean and fat flies then showed a net increase in fat percentage between 7 and 14 days, a typical response for insects with a surplus of food. Fat percentage remained high in lean females, but dropped in fat females, leading to females with low teneral fat reserves to become fatter at 21-28 days compared to females with high teneral fat reserves. The amber wasp *L. heterotoma* shows variation in the ability for adult fat accumulation and we hypothesized that variation to depend on genetic background. None of the inbred lines showed net fat accumulation, however, but the rate of fat depletion differed among inbred lines and depended on host developmental conditions. We further confirmed that *D. melanogaster* pupal size was highly correlated with adult female size for both *D. melanogaster* and *L. heterotoma.* Overall, our results for *D. melanogaster* show a typical age-response for fat accumulation, while our findings for *L. heterotoma* provide strong evidence that genetic background and developmental conditions shape the rate of fat depletion during adulthood.

**Summary statement:** Unlike their fruit fly hosts, parasitic wasps do not build up fat reserves as adults, and the speed at which they lose fat depends on their genetic background and host.

## 1. Introduction

Fat metabolism, or the synthesis and use of triacylglycerols, is a dynamic process (Arrese and Soulages, 2010; Scheifler et al., 2024). In animals, including insects, the physiological state, as well as the availability, amount, and composition of carbohydrates play a large role in determining the rate at which fat is synthesized and used. In *Drosophila melanogaster*, for example, larvae that were partially starved during development accumulated less fat compared to larvae that had continual access to sugar and yeast (Enriquez et al., 2022). Another study with *D. melanogaster* revealed that females only accumulate large triacylglycerol stores as adults when the developmental diet contains both sugar and yeast (Stefanyshyn et al., 2023). Insects will generally start accumulating fat for storage in the fat body when the diet contains a surplus of sugars and other carbohydrates (Arrese and Soulages, 2010; Visser et al., 2010; Visser and Ellers, 2008).

In many adult insects, accumulation of fat from sugars and other carbohydrates provides the energetic resources needed for maintenance and reproduction. In the cricket *Gryllus firmus,* artificially selected lines of flight-capable and flightless morphs differ in the rate at which triacylglycerols are synthesized (Zera, 2005). After 5 days of feeding, flight-capable morphs accumulated more fat to fuel dispersal. Flightless morphs did, however, still accumulate fat that was invested mainly in reproduction. Large fat stores were also deposited in the cricket *Gryllus bimaculatus,* where measures of fat content and tracing of triacylglycerol synthesis revealed an increase of triacylglycerols in early life that lowered as life progressed (Lorenz, 2001; Lorenz and Anand, 2004). The fat content of migrating monarch butterflies, *Danaus plexippus,* reaches a peak during the middle phase of their long-distance journey (Gibo and McCurdy, 1993). Early work with *D. melanogaster* revealed a steady increase of fat content as flies became older (Service, 1987 and see Box 1 in Scheifler et al., 2024) and similar results have been obtained also for other flies (Alqurashi et al., 2020). Taken together, these studies show that adults of many insects show a net increase in fat content from the first days to weeks after emergence when carbohydrates are available in the adult diet, whereas fat content often declines at older ages.

Parasitoids differ from other insects, because adults of most parasitoid species do not accumulate fat (Visser et al., 2010, Visser, LeLann et al., 2023; Visser and Ellers, 2008). Ellers (1996) already showed that fat content gradually decreases throughout adult life in the parasitoid *Asobara tabida.* The lack of fat accumulation (i.e., a net decrease in fat content) in *A. tabida,* but also other adult parasitoids, is not due to a lack of feeding (Ellers, 1996; Visser et al., 2010). Some parasitoid species remain capable of synthesizing small quantities of fatty acids (Giron & Casas, 2003; Ruther et al., 2021; Visser, LeLann et al., 2023). Feeding on a sugar-rich diet and even synthesis of some fatty acids does not, however, generally lead to the synthesis and accumulation of bulk fat as an energetic reserve (Visser et al., 2010; Visser, LeLann et al., 2023). A complete lack of fat accumulation is an exceptional adaptation that evolved as a consequence of the parasitoid lifestyle in many distinct parasitoid lineages (Visser et al., 2010).

The parasitoid amber wasp *Leptopilina heterotoma* is a koinobiont endoparasitoid of *Drosophila* larvae, including *D. melanogaster* (Quicray et al., 2023). Female parasitoids oviposit in second-to third-instar host larvae, where the wasps continue to feed until pupation. The parasitoid larva consumes all host tissues, including the host’s fat body, before emerging as an adult from the host puparium (Quicray et al., 2023). The host thus represents the entire developmental environment of the wasp, and fat reserves at adult emergence (i.e., the teneral reserves) are thus mainly host-derived. The first empirical data on *L. heterotoma* revealed that fat reserves were accumulated in adults (i.e., a net increase in fat content; Eijs et al., 1998; Visser et al., 2010). A large-scale study comparing fat accumulation and use between different *Leptopilina* species then revealed that none of the species, including *L. heterotoma,* accumulated fat (Visser et al., 2018). Testing various populations on different host strains then revealed that wasps kept on a fatter *D. melanogaster* strain did not accumulate fat (2016 populations in Visser et al., 2018), while populations kept on a leaner *D. melanogaster* strain did accumulate fat (2013 populations in Visser et al., 2018). Further tests with different host species (i.e., *D. melanogaster* and *Drosophila simulans*), and hosts that differed in fat content (lean and fat *D. melanogaster*) revealed that fat accumulation of field-caught *L. heterotoma* populations also varied (Visser et al., 2021). In that study, host fat content was measured directly, and stable-isotope tracing demonstrated that *L. heterotoma* fatty acid synthesis depended on fat content of the hosts, i.e., that fatty acid synthesis is phenotypically plastic (Visser et al., 2021). For fat accumulation, previous results suggest that there is population-level variability, where fat content of the host plays a key role in determining wasp phenotypes, but we have yet to understand the role played by the genotype of the wasp.

Here, we define plasticity of fat accumulation as variation in the fat-content trajectory of a given wasp genotype across developmental host conditions (complementing previous work on plasticity of fatty acid synthesis *in L. heterotoma*; Visser et al., 2021). We test how age-dependent changes in whole-body fat content during adulthood differ between two insect species, *D. melanogaster* and its parasitoid wasp *L. heterotoma*. To manipulate the amount of fat available at adult emergence, we generated lean and fat *D. melanogaster* by varying larval density and food amount (as in Enriquez et al., 2022). Fat content of lean and fat adult flies was then measured throughout adulthood. We hypothesized that our *D. melanogaster* strain would show a typical insect response where fat is accumulated irrespective of the amount of fat at emergence. We expected that fat flies would accumulate less fat compared to lean flies. Lean and fat *D. melanogaster* larvae also served as developmental hosts for *L. heterotoma*. We generated 5 *L. heterotoma* inbred lines to uncover genotype-level variation. We hypothesized that substantial fat accumulation would take place when wasps developed in lean hosts, but not in fat hosts. We expected most fat to be accumulated during the first weeks of adult female life, a time at which most fat is needed as energetic fuel in nature. We further expected to see different patterns of fat accumulation depending on the genotype. We also collected data on *D. melanogaster* pupal size and body size of *D. melanogaster* and *L. heterotoma* to show that the size of the empty pupal casing provides a proxy of the host fat resources available to the developing wasp. To the best of our knowledge, this is the first study to characterize age-dependent fat-content trajectories throughout adult life (up to 56 days after emergence) in a parasitoid and non-parasitoid insect reared under identical conditions. By using 5 distinct inbred *L. heterotoma* lines, we investigated for the first time how age-dependent fat-content trajectories depend on genetic background and developmental host conditions (genotype-by-environment-by-age interaction).

## 2. Materials and methods

### 2.1. Insects

#### 2.1.1. The vinegar fly (host) *D. melanogaster*

The vinegar fly *D. melanogaster* was kindly provided by Patricia Gibert (ULyon, France). This strain was originally obtained from field collections in 1994 in Sainte-Foy-les-Lyon and maintained in our laboratory since 2016. For experiments, we generated lean and fat *D. melanogaster* by manipulating both larval density and nutrient content. Crowding reduces food availability for developing individuals leading to lower fat reserves (Enriquez et al., 2022; Henry et al., 2018). To generate lean larvae, adults were allowed to lay eggs for 15-16 hours, after which 300 eggs were placed in 1mL of standard medium (consisting of 20 g agar, 35 g yeast, 50 g sugar, 10 ml nipagin, and 5 ml propionic acid per liter water). To generate fat larvae, 150 eggs were placed in 5mL of sugar-rich medium (containing twice the amount of sugar in 1 liter water compared to the standard medium). Lean and fat *D. melanogaster* pupae were sexed and female pupae placed singly in a small plastic vial (5 mL). Upon emergence, females were either immediately frozen or allowed to feed on the standard medium for 7, 14, 21, 28, 35, 42, 49 or 56 days, after which females were frozen. Food was available *ad libitum* and renewed every week. Females were then used to measure triacylglycerol content using gravimetry (see below). Sample sizes (n females for developmental condition × age) are reported in Table 1. Experiments were performed at 23°C, 75% relative humidity and a photoperiod of L:D 16:8, mimicking summer conditions in temperate Europe and allowing good development and survival of *D. melanogaster* and *L. heterotoma*. These protocols produce two non-overlapping size categories in our *Drosophila* strain (Enriquez et al., 2022), which we verified by measuring the pupal size of every female (see results). Females (and their associated pupa) were collected at random within each treatment. Adult age was counted from the day of emergence of each female (emergences were checked daily), so that the 15-16 h egg-laying window had no effect on age at sampling. Our aim was to compare two contrasting developmental conditions, so no intermediate control group was included.

**Table 1.** Fat percentage (mean ± SE) during adulthood (0-56 days after emergence) for adult female *D. melanogaster* reared under fat and lean developmental conditions, and for adult females of five *L. heterotoma* inbred lines reared on fat and lean *D. melanogaster* hosts. n = number of females (each measured once) per *D. melanogaster*/*L. heterotoma* inbred line x developmental condition x age (in days). Letters indicate significant fat percentage differences between ages within each species (or inbred line) x developmental condition (post hoc pairwise comparisons of estimated marginal means with Tukey adjustment, p < 0.05). Empty cells indicate that no living females remained at that age.

|  | Fat percentage |  |  |  |
|---|---|---|---|---|
|  | Fat |  | Lean |  |
| Age | n | Mean ± 1 SE (%) | n | Mean ± 1 SE (%) |
| *D. melanogaster* |  |  |  |  |
| 0 | 30 | 21.94 ± 0.74<sup>a</sup> | 32 | 10.71 ± 0.67<sup>ab</sup> |
| 7 | 30 | 15.24 ± 0.84<sup>b</sup> | 31 | 9.61 ± 0.66<sup>a</sup> |
| 14 | 31 | 19.45 ± 0.97<sup>ac</sup> | 34 | 22.05 ± 1.12<sup>c</sup> |
| 21 | 30 | 12.45 ± 0.65<sup>b</sup> | 31 | 20.91 ± 1.05<sup>c</sup> |
| 28 | 30 | 14.49 ± 0.93<sup>b</sup> | 33 | 20.98 ± 1.08<sup>c</sup> |
| 35 | 31 | 15.53 ± 1.28<sup>bc</sup> | 30 | 17.85 ± 1.16<sup>cd</sup> |
| 42 | 31 | 13.73 ± 1.05<sup>b</sup> | 36 | 17.59 ± 1.11<sup>cd</sup> |
| 49 | 34 | 15.3 ± 1.27<sup>bc</sup> | 31 | 15.07 ± 1.17<sup>d</sup> |
| 56 | 30 | 14.75 ± 1.18<sup>bc</sup> | 21 | 15.27 ± 2.26<sup>bcd</sup> |
| *L. heterotoma* I1 |  |  |  |  |
| 0 | 30 | 33.03 ± 0.76<sup>a</sup> | 28 | 18.26 ± 1.39<sup>a</sup> |
| 7 | 31 | 16.1 ± 0.42<sup>b</sup> | 15 | 14.33 ± 0.79<sup>ab</sup> |
| 14 | 17 | 12.52 ± 0.91<sup>cd</sup> | 11 | 11.86 ± 0.58<sup>b</sup> |
| 21 | 17 | 14.86 ± 0.96<sup>bc</sup> | 10 | 11.64 ± 1.02<sup>b</sup> |
| 28 | 17 | 10.76 ± 0.79<sup>d</sup> | 6 | 9.22 ± 1.64<sup>b</sup> |
| 35 | 17 | 10.41 ± 0.58<sup>d</sup> |  |  |
| 42 | 17 | 10.21 ± 0.59<sup>d</sup> |  |  |
| 49 | 5 | 9.2 ± 0.75<sup>cd</sup> |  |  |
| *L. heterotoma* I5 |  |  |  |  |
| 0 | 30 | 34.36 ± 0.72<sup>a</sup> | 30 | 20.06 ± 1.21<sup>a</sup> |
| 7 | 30 | 19.17 ± 0.66<sup>b</sup> | 15 | 11.37 ± 1.07<sup>b</sup> |
| 14 | 17 | 15.23 ± 0.76<sup>c</sup> | 10 | 12.91 ± 0.89<sup>b</sup> |
| 21 | 17 | 13.81 ± 0.81<sup>cd</sup> | 2 | 11.15 ± 0.81<sup>b</sup> |
| 28 | 17 | 11.55 ± 0.43<sup>de</sup> |  |  |
| 35 | 17 | 10.69 ± 0.54<sup>e</sup> |  |  |
| 42 | 6 | 9.45 ± 0.64<sup>e</sup> |  |  |
| *L. heterotoma* I7 |  |  |  |  |
| 0 | 30 | 31.56 ± 0.74<sup>a</sup> | 32 | 17.43 ± 1.33<sup>a</sup> |
| 7 | 30 | 18.71 ± 0.58<sup>b</sup> | 15 | 14.9 ± 1.68<sup>ab</sup> |
| 14 | 17 | 17.58 ± 0.7<sup>bc</sup> | 10 | 14.25 ± 0.62<sup>a</sup> |
| 21 | 17 | 15.93 ± 0.7<sup>bc</sup> | 10 | 10.12 ± 0.64<sup>b</sup> |
| 28 | 17 | 14.13 ± 0.93<sup>cd</sup> | 3 | 8.54 ± 2.46<sup>ab</sup> |
| 35 | 17 | 11.88 ± 0.96<sup>de</sup> |  |  |
| 42 | 19 | 11.11 ± 0.73<sup>de</sup> |  |  |
| 49 | 17 | 7.47 ± 0.58<sup>f</sup> |  |  |
| 56 | 4 | 6.73 ± 1.38<sup>ef</sup> |  |  |
| *L. heterotoma* I8 |  |  |  |  |
| 0 | 30 | 32.42 ± 0.55<sup>a</sup> | 19 | 19.7 ± 1.83<sup>a</sup> |
| 7 | 30 | 19.33 ± 0.88<sup>b</sup> | 15 | 16.14 ± 0.74<sup>ab</sup> |
| 14 | 17 | 18.33 ± 0.88<sup>b</sup> | 10 | 12.42 ± 1.21<sup>b</sup> |
| 21 | 17 | 17.55 ± 0.93<sup>b</sup> | 10 | 12.83 ± 1.32<sup>ab</sup> |
| 28 | 17 | 14.98 ± 0.87<sup>bc</sup> |  |  |
| 35 | 17 | 12.68 ± 1.18<sup>c</sup> |  |  |
| 42 | 18 | 12.34 ± 1.31<sup>c</sup> |  |  |
| 49 | 14 | 11.3 ± 1.56<sup>c</sup> |  |  |
| *L. heterotoma* I10 |  |  |  |  |
| 0 | 30 | 31.63 ± 0.92<sup>a</sup> | 23 | 16.66 ± 1.59<sup>a</sup> |
| 7 | 30 | 16.85 ± 0.84<sup>b</sup> | 15 | 13.97 ± 1.26<sup>a</sup> |
| 14 | 17 | 16.77 ± 0.94<sup>b</sup> | 12 | 14.79 ± 0.95<sup>a</sup> |
| 21 | 17 | 14.95 ± 0.77<sup>bc</sup> | 10 | 14.58 ± 1.9<sup>a</sup> |
| 28 | 17 | 11.72 ± 1.1<sup>cd</sup> | 3 | 9.88 ± 2.27<sup>a</sup> |
| 35 | 17 | 9.89 ± 0.63<sup>d</sup> |  |  |
| 42 | 17 | 9.62 ± 0.57<sup>d</sup> |  |  |
| 49 | 3 | 7.9 ± 0.21<sup>bcd</sup> |  |  |

#### 2.1.2. The amber wasp *L. heterotoma*

We used 5 inbred lines generated from a population that was originally collected in Eupen, Belgium, in 2016. Lines were inbred through sister-brother matings and used after >40 generations. For experiments, lean and fat *D. melanogaster* larvae (generated as described above) were harvested 2 days after egg collection. At this time, *D. melanogaster* larvae are in the second to third larval instar, which is ideal for *L. heterotoma* development (Quicray et al., 2023). One newly emerged *L. heterotoma* female and two males of each inbred line were allowed to mate, after which females were allowed to lay eggs either in lean or fat larvae until death of the female. As the host represents the entire developmental environment of the wasp, and we are interested in variation in host fat content (Enriquez et al., 2022), we maintained each parasitized pupa in a separate tube until adult wasp emergence. Upon emergence, females were either frozen or allowed to feed on a 2:1 water:honey solution during 7, 14, 21, 28, 35, 42, 49 or 56 days, after which females were frozen. Females were then used to measure triacylglycerol content using gravimetry. As for *D. melanogaster*, parasitized pupae were collected at random. Experimental conditions were similar to those described above for *D. melanogaster.* Sample sizes (n females for each genotype × developmental host condition × age) are reported in Table 1.

### 2.2. Experimental set-up: gravimetric method to determine fat content

Fat content was determined using standardized gravimetric methods, as described in (Enriquez et al., 2022; Visser et al., 2010, 2018). Briefly, flies and wasps were dried for 3 days at 60℃, after which dry weight was determined on an ultra-precise balance (Cubis 3.6P, Sartorius AG, Göttingen, Germany). Insects were then placed in 4mL diethyl ether for 24 hours. Insects were rinsed with 1mL diethyl ether and dried again for 3 days at 60℃. Triacylglycerol content was then determined by subtracting dry weight after ether extraction from dry weight before ether extraction. We calculated the percentage of fat based on the absolute amount of triacylglycerol and the dry weight before extraction.

### 2.3. Pupal and tibia size measurements

Pupal size is a validated proxy for fat content in non-parasitized pupae of the *D. melanogaster* strain used here (Enriquez et al., 2022). As the developing wasp feeds exclusively on its host, the fat reserves of a newly emerged wasp reflect the resources available in the host. The size of the empty host pupal casing was, therefore, used as a proxy of fat availability. Tibia length is a common indicator of fly and wasp body size (Ellers, 1996; Ellers et al., 1998; Enriquez et al., 2022). To measure *D. melanogaster* pupal size (or, for wasps, of the empty host pupal casing remaining after wasp emergence), a picture of each pupa was taken using a Zeiss microscope (Stemi 508, Axocam 208 color) and the total area of the pupal case (mm^2^) measured using ImageJ software (v1.54f) (Schneider et al., 2012). To measure adult fly and wasp tibia length, the right hind leg was removed, photographed, and the proximal to distal length (µm) estimated, also using ImageJ software.

### 2.4. Statistics

All analyses were conducted with R project (v4.6.1, R Core Team, 2026). Statistical significance was set at p < 0.05. Age-dependent changes in fat percentage during adulthood for both *D. melanogaster* and *L. heterotoma* were analyzed using generalized linear models (GLMs). The distribution of the data was assessed using goodness-of-fit criteria (AIC/BIC) with the *fitdistrplus* package (Delignette-Muller and Dutang, 2015), and a Gamma (log link) distribution was selected for *D. melanogaster* and a log-normal distribution for *L. heterotoma*. Fly and wasp females, each measured once, were the experimental unit (biological replicate) in all analyses. Females were sampled at random at each age.

For *D. melanogaster*, to test whether age-dependent changes in fat percentage differed between phenotypes, the full GLM included fly phenotype (fat vs. lean), age (days 0-56, categorical factor), as well as their interaction. As dispersion differed among phenotype x age combinations in *D. melanogaster* (Levene’s test on Pearson residuals, p < 0.01), the Gamma GLM was fitted with a dispersion parameter specific to each phenotype × age combination (*glmmTMB* package; Brooks et al., 2017), which improved model fit compared with a constant dispersion (likelihood-ratio test, p < 0.001). Model significance was evaluated using type III Wald chi-square tests (Anova function, *car* package; Fox & Weisberg, 2019). Post hoc comparisons were performed on the full model using estimated marginal means (*emmeans* package, Lenth & Piaskowski, 2025): fat percentage was compared between ages within each phenotype with Tukey adjustment, and between phenotypes at each age with Holm adjustment.

For *L. heterotoma*, the full model included host phenotype (fat vs. lean *D. melanogaster*), age (days 0-56, categorical factor), inbred line (I1, I5, I7, I8, and I10), as well as all interactions, and was fitted to log-transformed fat percentage with a Gaussian distribution (i.e., a log-normal model). As variance differed among groups (Levene’s test, p < 0.001), several dispersion structures varying with age, host phenotype and inbred line were compared using AIC and BIC (*glmmTMB* package). The selected model included a dispersion parameter specific to each host phenotype × age combination and an effect of inbred line on dispersion, which strongly improved model fit compared with a constant dispersion (likelihood-ratio test, p < 0.001). A model with a dispersion parameter for each inbred line × host phenotype × age combination was not retained, as it was strongly penalized by BIC and estimated dispersion from groups of only two to three females. Parameters that could not be estimated because some host phenotype × age combinations were missing at older ages were dropped from the model. Model significance was evaluated using type II Wald chi-square tests (Anova function, *car* package), as type III tests are not appropriate when some combinations are missing. Post hoc comparisons were performed on the full model using estimated marginal means (*emmeans* package): fat percentage was compared between ages within each inbred line × host phenotype with Tukey adjustment.

Differences in pupal size and adult female tibia length between fly phenotypes were analyzed using Wilcoxon rank-sum tests for *D. melanogaster*. For *L. heterotoma*, linear models were used to account for differences in genetic background between inbred lines. Pupal size and adult female tibia length were each modeled as a function of inbred line, *D. melanogaster* host phenotype, and their interaction. Model significance for *L. heterotoma* was assessed using type II or type III ANOVA, depending on the significance of the interaction, and host phenotypes were then compared within each inbred line using estimated marginal means with Holm adjustment. Normality of residuals and homogeneity of variances were checked with Shapiro-Wilk and Levene’s tests; variances differed only moderately among groups. The proportion of variance explained by each factor was estimated with eta-squared (*effectsize* package; Ben-Shachar et al., 2020).

## 3. Results

### **3.1.** Age-dependent changes in fat content during adulthood in D. melanogaster

We first examined how fat content changed with age in adult *D. melanogaster* reared under lean and fat developmental conditions, as a non-parasitoid reference. Age-dependent changes in fat percentage differed between lean and fat *D. melanogaster* phenotypes (age × phenotype interaction: p < 0.001; Table 1; Figure 1). Fat percentages changed significantly with age (p < 0.001) but did not differ between *D. melanogaster* phenotypes when averaged across all ages (p > 0.05), because the trajectories of lean and fat flies crossed during adulthood. Lean flies emerged with a fat percentage of 10.71 ± 0.67% (mean ± 1 SE), which did not change significantly during the first week of adult life (9.61 ± 0.66% at 7 days; p > 0.05). Fat percentage then increased steeply to a peak of 22.05 ± 1.12% at 14 days and gradually declined with age, being significantly lower at 49 days than at 14 days (15.07 ± 1.17%; p < 0.05). At 56 days, fat percentage (15.27 ± 2.26%) was highly variable and no longer differed significantly from either the peak at 14 days or emergence. A different pattern was observed for fat flies that emerged with 21.94 ± 0.74% of fat, which decreased by 6.7 percentage points during the first 7 days of adult life (to 15.24 ± 0.84%, p < 0.05). Fat percentage then increased significantly to 19.45 ± 0.97% at 14 days (p < 0.05), a value similar to that at emergence, before dropping significantly to 12.45 ± 0.65% at 21 days (p < 0.05; Table 1). It did not change significantly thereafter, ranging between 12.45% and 15.53% until 56 days, when it reached 14.75 ± 1.18%, a value similar to that of lean flies. Fat percentage of fat flies remained lower than at emergence at all ages except 14 days.

**Figure 1.**
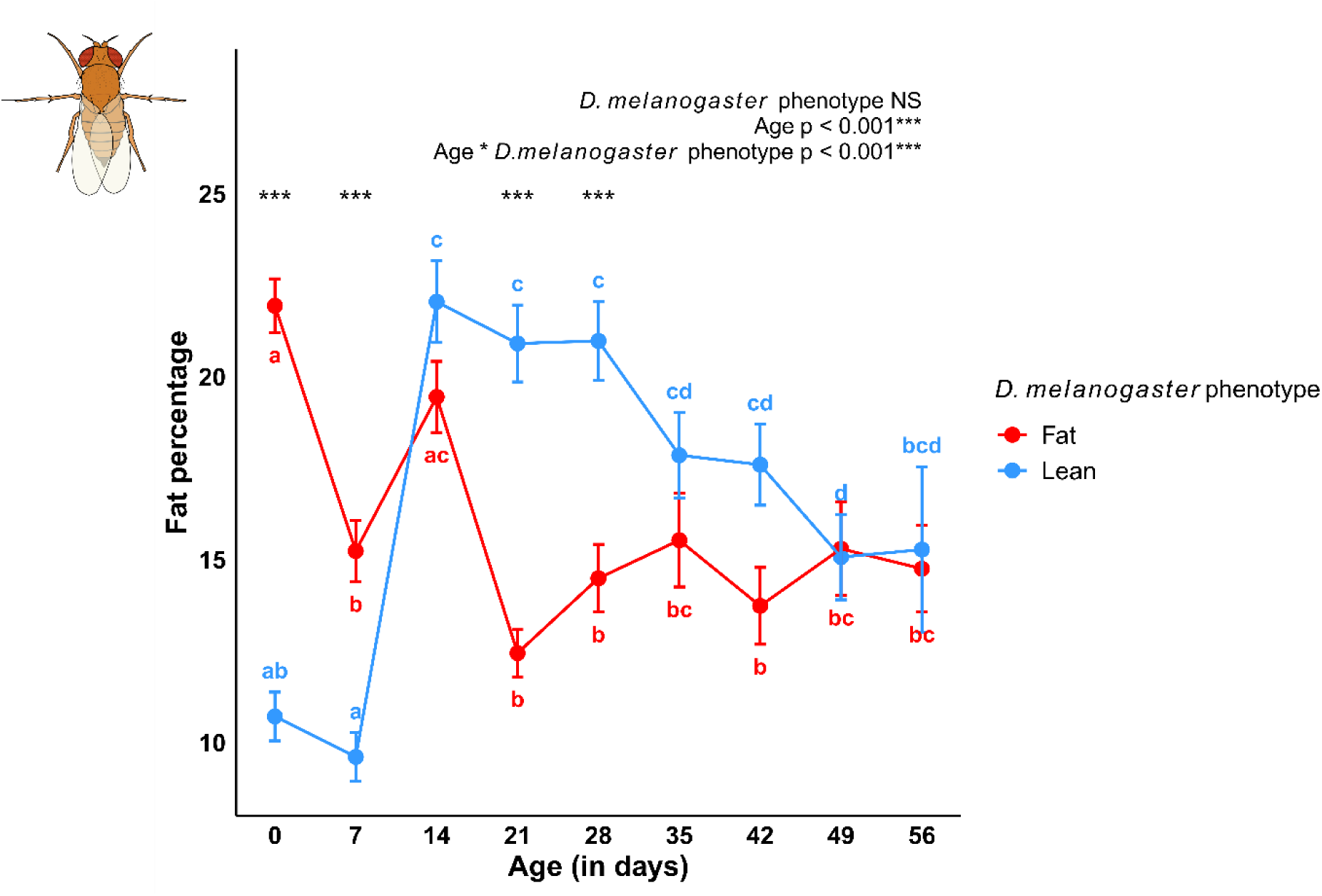
Age-dependent changes in fat percentage during adulthood in female *D. melanogaster* reared under fat (red) and lean (blue) developmental conditions. Fat percentage was measured at 0, 7, 14, 21, 28, 35, 42, 49 and 56 days after emergence; each female was measured once. Points represent means and error bars ± 1 SE; n = 21-36 females per developmental condition × age (Table 1). Results of the Gamma GLM with group-specific dispersion (type III Wald tests) testing the effects of developmental condition, age and their interaction on fat percentage are reported at the top of the graph. Different letters indicate significant differences between ages within each developmental condition (post hoc comparisons of estimated marginal means with Tukey adjustment, p < 0.05). Asterisks indicate significant differences between fat and lean flies at a given age (estimated marginal means with Holm adjustment; ***: p < 0.001). To increase readability, the y-axis does not start at 0.

Pairwise comparisons between *D. melanogaster* phenotypes at each age showed that fat flies had a higher fat percentage than lean flies at emergence (∼2 times higher) and at 7 days (∼1.6 times higher; p < 0.001 for both). In absolute terms, fat flies emerged with about nine times more fat than lean flies (64.5 ± 3.0 µg vs. 7.0 ± 0.5 µg; Welch’s t-test, p < 0.001). This pattern was reversed later in adult life, with lean flies having a higher fat percentage than fat flies at 21 days (∼1.7 times higher; p < 0.001) and 28 days (∼1.4 times higher; p < 0.001). Fat percentage did not differ between phenotypes at 14, 35, 42, 49 and 56 days (p > 0.05; Figure 1).

### **3.2.** Pupal and adult female size in D. melanogaster

To verify that the two developmental treatments produced distinct host phenotypes, we measured pupal size and adult female tibia length. Lean and fat *D. melanogaster* differed strongly in both pupal size and adult female size (measured as tibia length), with no overlap between the size distributions of lean and fat females (p < 0.001; Figure 2). Lean pupae were on average 1.30 ± 0.009 mm² (mean ± 1 SE), while fat pupae were on average 2.77 ± 0.012 mm² (Figure 2). Fat pupae were thus more than twice as big as lean pupae. For adult female size, tibia length was on average 511 ± 1.98 μm for lean females, while tibia length of fat females was on average 727 ± 1.36 μm (Figure 2). Fat females were thus almost 1.5 times bigger than lean females.

**Figure 2.**
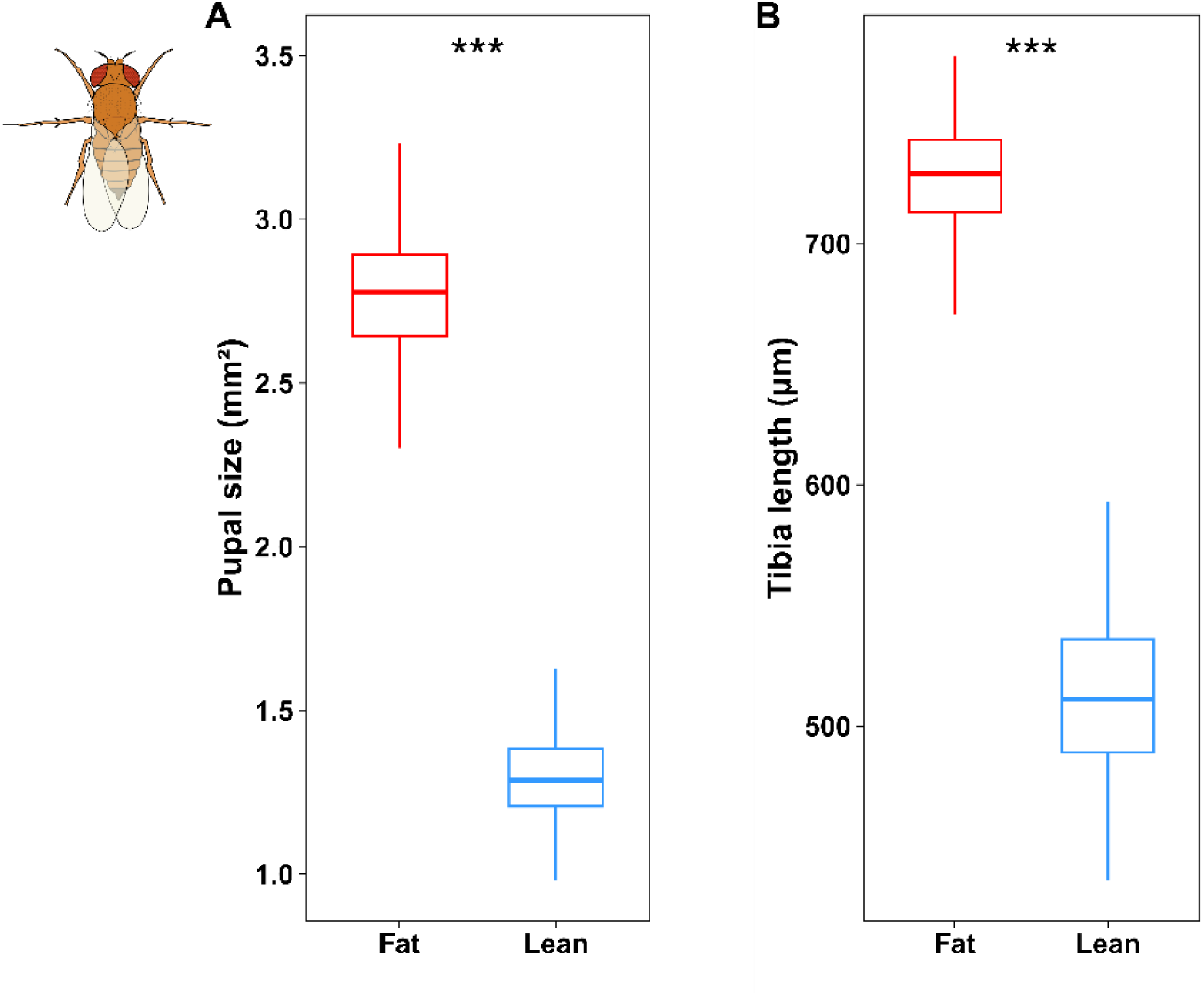
(A) Pupal size (mm²) and (B) adult female tibia length (μm) of *D. melanogaster* reared under fat (red) and lean (blue) developmental conditions. Boxplots show medians and interquartile ranges for each developmental condition (fat: n = 277, lean: n = 279 females). Pupal size and tibia length differences between developmental conditions were tested with Wilcoxon rank-sum tests (***: p < 0.001). To increase readability, the y axis does not start at 0.

### **3.3.** Age-dependent changes in fat content during adulthood in inbred lines of L. heterotoma

We then examined whether age-dependent changes in fat content of adult *L. heterotoma* depended on inbred line and on the developmental host condition. Age-dependent changes in fat percentage during adulthood differed between inbred lines and between *L. heterotoma* reared on lean or fat *D. melanogaster* (i.e., host phenotype; p < 0.05 for the three-way interaction between inbred line, host phenotype, and age; Table 1; Figure 3). Wasps of all inbred lines reared on fat *D. melanogaster* emerged with a very high average percentage of fat, 32.6 ± 0.34% (mean ± 1 SE). Fat percentage decreased by almost 15% during the first 7 days of adult life (to 18.0% on average across inbred lines) and continued to decline with increasing age (no significant increase at any age in any inbred line; Table 1). Wasps reared on lean hosts emerged with a considerably lower fat percentage, on average 18.4 ± 0.64%, which did not increase thereafter. Fat percentage decreased significantly during the first week of adult life only in line I5. At later ages, fat percentage was also significantly lower than at emergence in lines I1 (14, 21 and 28 days), I7 (21 days) and I8 (14 days), whereas it did not change significantly with age in line I10 (Table 1). No wasps reared on lean hosts were available for sampling after 21 (I5, I8) or 28 days (I1, I7, I10) of adult life, whereas wasps reared on fat hosts were sampled until 42-56 days, depending on the inbred line (Table 1). Inbred lines, representing different genotypes, differed significantly in the rate at which fat percentage declined with age (age x inbred line, p < 0.001; Table 1; Figure 3). For example, in wasps reared in fat hosts, fat percentage at 49 days ranged from 7.47 ± 0.58% in the inbred line I7 to 11.30 ± 1.56% in the inbred line I8 (Table 1).

**Figure 3.**
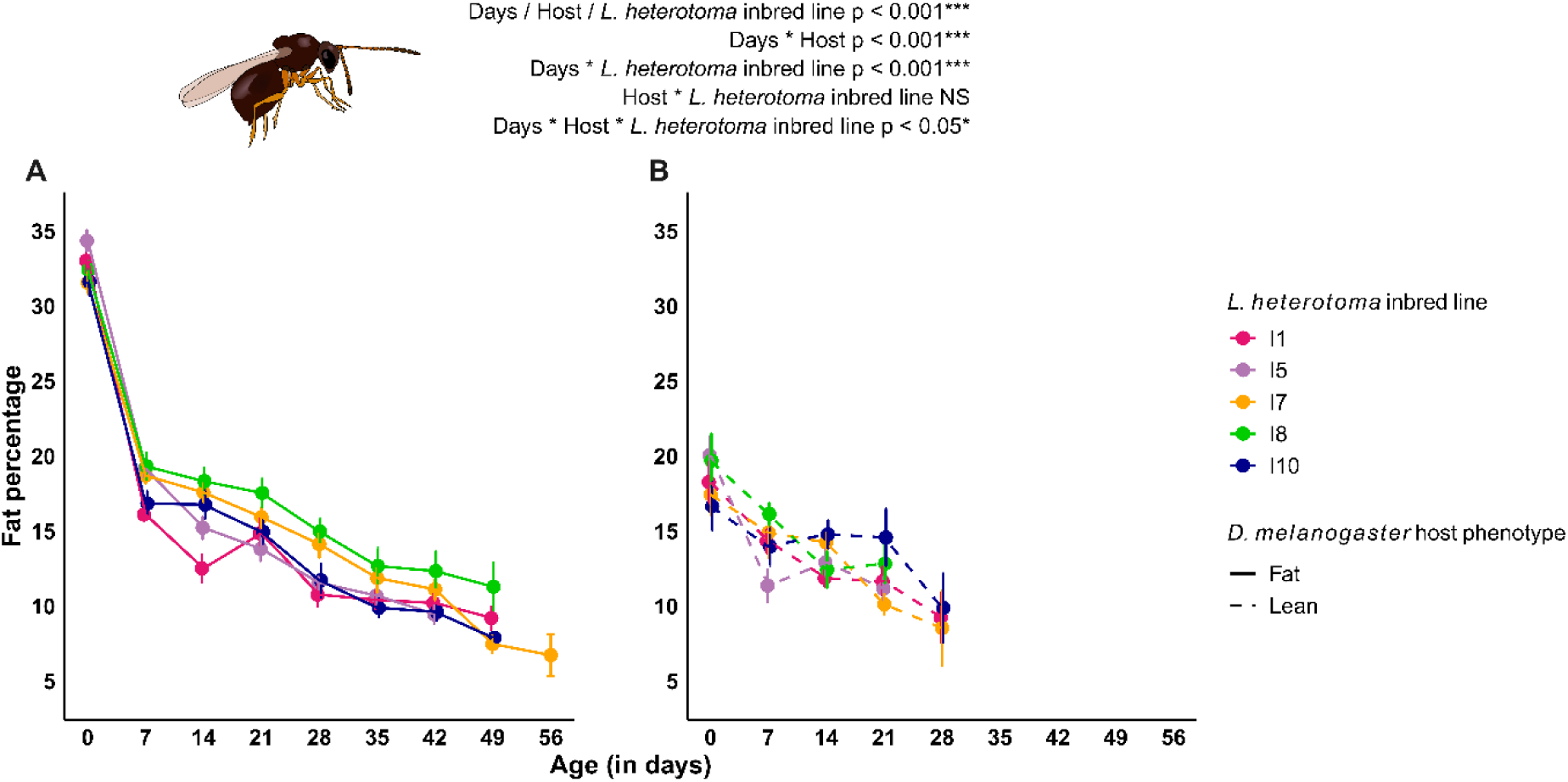
Age-dependent changes in fat percentage during adulthood in females of five *L. heterotoma* inbred lines (I1, I5, I7, I8 and I10) reared on (A) fat and (B) lean *D. melanogaster* hosts. Fat percentage was measured at 0, 7, 14, 21, 28, 35, 42, 49 and 56 days after emergence; each female was measured once. Points represent means and error bars ± 1 SE; n = 2-32 females per inbred line x developmental condition × age (Table 1). Wasps reared on lean hosts were not available (absence of living females) for sampling after 21 (I5, I8) or 28 days (I1, I7, I10), and wasps reared on fat hosts after 42 (I5), 49 (I1, I8, I10) or 56 days (I7). Results of the log-normal model with group-specific dispersion (type II Wald tests) testing the effects of inbred line, developmental condition, age and all interactions on fat percentage are reported at the top of the graph. Differences between ages within each inbred line x developmental condition (post hoc comparisons of estimated marginal means with Tukey adjustment, p < 0.05) are reported in Table 1. Inbred lines are shown with the same colors in both panels, which share the same y-axis scale. To increase readability, the y-axis does not start at 0.

### **3.4.** Host pupal size and adult female size in L. heterotoma

To evaluate how developmental host conditions affected wasp size, we measured the empty host pupal casing and adult female wasp tibia length. For all inbred lines, host pupal sizes differed ∼1.8 times between wasps reared on lean and fat *D. melanogaster* phenotypes (p < 0.001; Figure 4), with an average of 1.38 ± 0.009 mm² (mean ± 1 SE) for wasps reared on lean hosts and an average of 2.52 ± 0.008 mm² for wasps reared on fat hosts. Adult female wasps reared on lean hosts were also smaller (tibia length ∼1.4 times shorter), with an average size of 426 ± 1.86 μm, compared to wasps reared on fat hosts with an average size of 592 ± 1.05 μm (Figure 4).

**Figure 4.**
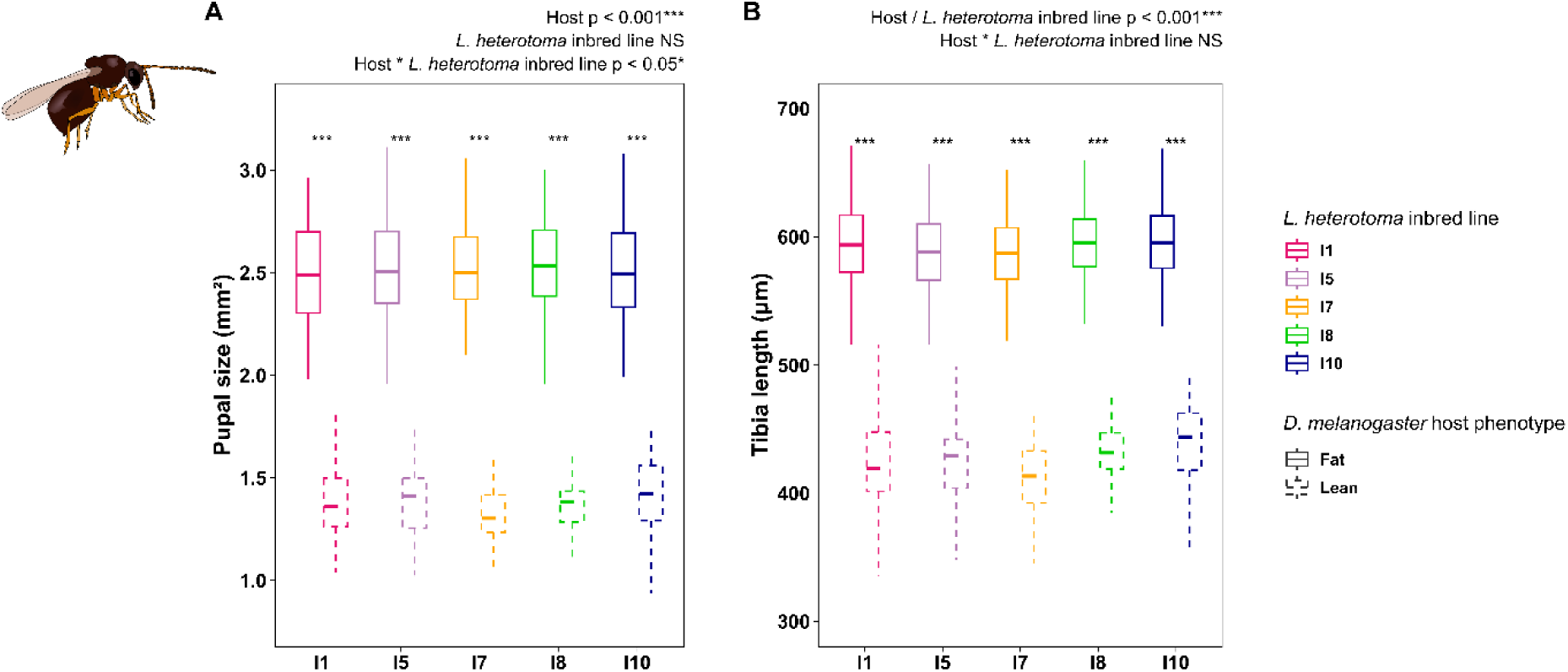
(A) Pupal size (mm²) and (B) adult female tibia length (μm) of five *L. heterotoma* inbred lines reared on fat (solid lines) and lean (dashed lines) *D. melanogaster* hosts. Boxplots show medians and interquartile ranges for each inbred line × developmental condition; n = fat/lean females: I1 n=151/70; I5 n=134/57; I7 n=168/70; I8 n=160/54; I10 n= 148/63). Results of the linear models testing the effects of inbred line, host phenotype and their interaction are reported at the top of each panel, and asterisks on the graphs indicate significant differences between developmental host conditions within each inbred line (post hoc comparisons of estimated marginal means with Holm adjustment; ***: p < 0.001). To increase readability, the y axis does not start at 0.

Both pupal size and tibia length differed strongly between developmental conditions in all inbred lines (p < 0.001), with host phenotype explaining most of the variance (η² = 0.85 and 0.86, respectively), whereas inbred line and the interaction between inbred line and host phenotype each explained less than 1% of the variance. Inbred line had no effect on pupal size (p > 0.05) and only a minor, although significant, effect on tibia length (p < 0.001). The difference between host conditions in pupal size varied slightly among inbred lines (inbred line × host phenotype interaction, p = 0.03; from 1.09 mm² in I1 and I10 to 1.19 mm² in I7), whereas no such interaction was found for tibia length (p > 0.05).

## 4. Discussion

Age-dependent patterns of fat accumulation and depletion during adulthood were very different for the vinegar fly *D. melanogaster* and inbred lines of the parasitoid amber wasp *L. heterotoma.* In *D. melanogaster*, the fat content of lean females remained stable during the first week of adult life and then increased by 12.4%. Fat females, in contrast, lost fat during the first week, before a significant but transient increase at 14 days. Net fat accumulation was thus observed in both phenotypes (although it was much stronger in lean females), in line with our expectations for an insect with typical fat accumulation. From 35 days onwards, fat percentages no longer differed between lean and fat flies, which converged to same values at 49 and 56 days, a considerable age for small insects. Despite the major disadvantage of having only little fat reserves for lean flies at emergence, lean females compensate for lower initial reserves through a net accumulation of fat during early adulthood. In contrast to our expectations for *L. heterotoma,* no fat accumulation was detected in any of the inbred lines during adulthood, irrespective of whether the wasps developed on lean or fat *D. melanogaster* hosts. Wasps reared on fat hosts thus emerged with much more fat reserves, which is consistent with the reduced apparent survival of lean wasps: no more lean wasps were available for sampling after 28 days of adult life, whereas fat wasps were sampled until 42-56 days of age (depending on *L. heterotoma* inbred line). We further showed that large differences can be generated in *D. melanogaster* pupal size, leading to large differences in both fly and wasp adult female size.

### **4.1.** Manipulating D. melanogaster size beyond natural phenotypic values

We generated extreme size differences between lean and fat *D. melanogaster*, both at the pupal and adult stage. Large-size differences result from the well-documented developmental plasticity of size for *D. melanogaster* (Enriquez et al., 2022; Shingleton et al., 2009). Larval density is one of the key environmental drivers of size-related plasticity: as density increases, food availability per larva decreases, leading to smaller pupae and consequently smaller adults. Nutritional quality during larval development also plays an important role, as starvation and reduced nutrient content further reduce pupal size (Enriquez et al., 2022). Our lean and fat *D. melanogaster* pupae (average 1.30 mm² and 2.77 mm², respectively) were within the range observed in a laboratory experiment performed by Enriquez et al. (2022), who generated *D. melanogaster* individuals of varying sizes (by manipulating both larval density and nutritional conditions) using the same *D. melanogaster* strain. Enriquez et al. (2022) also measured pupal size of field-caught individuals across five *Drosophila* species, including *D. melanogaster*, and found that none fell below approximately 1.5 mm². Our laboratory-reared lean females thus represent an extreme that is not common in nature, while generated fat flies are comparable to the range of sizes observed in natural populations.

In insects, pupal size is widely used as a proxy for body size and fat content (Enriquez et al., 2022; Koenraadt, 2008). In *D. melanogaster*, larger pupae consistently have a higher fat content, both in laboratory-reared and field-caught individuals. Field-caught individuals are, however, generally larger and fatter than laboratory individuals (Enriquez et al., 2022). The strong positive relationship between pupal size and adult female tibia length we observed here is, therefore, expected: both traits reflect the same variation in body size driven by larval development. Although Enriquez et al. (2022) did not measure adult tibia length, our values are consistent with those reported in the literature for laboratory-reared *D. melanogaster* (Siomava et al., 2016), with fat females (average tibia length of 727 μm) being considerably larger than lean females (average tibia length of 511 μm), mirroring the pattern observed at the pupal stage.

For *L. heterotoma*, the large differences between wasps reared on lean and fat hosts (on average 426 μm versus 592 μm for female tibia length) are in line with the well-established role of host size in determining parasitoid adult body size (Cohen et al., 2005). In our study, host phenotype explained 85-86% of the variance in *L. heterotoma* pupal size and tibia length, whereas inbred line explained less than 1%, confirming that adult wasp size was mostly determined by developmental host condition rather than genetic background. Wang et al. (2021), for example, showed that wasp body size was positively correlated with host size considering 25 drosophilid host species and two parasitoid species, *Pachycrepoideus vindemiae* and *Trichopria drosophilae*. Our lean *L. heterotoma* females that developed in hosts smaller than what is typically found in nature (Enriquez et al., 2022), therefore, represent a size class that is below the lower threshold of what is commonly encountered in the field. The very small size of lean wasps could explain why no net fat accumulation was detected, possibly as a consequence of a physical rather than a metabolic constraint. Wasps with such small body sizes may lack the physical space and body storage capacity needed for additional fat accumulation. This hypothesis remains to be tested, as we did not measure lipid synthesis directly. It is, however, supported by the strongly reduced apparent survival of lean wasps, with no females remaining alive after 28 days compared to 42-56 days for fat wasps, suggesting the limit of viability for lean females. Future studies comparing *L. heterotoma* derived from hosts with more natural sizes could determine if the absence of fat accumulation in lean inbred wasps is indeed a physical constraint or a plastic response that can still be reversed under more natural developmental conditions (Visser et al., 2018, 2021).

### 4.2. *Patterns of fat accumulation and depletion in* Drosophila

Recent years have seen an increase in studies on fat-related traits in *D. melanogaster* (Heier and Kühnlein, 2018; Schlame et al., 2020; Trinh and Boulianne, 2013). While the increase in knowledge on fat metabolism in *D. melanogaster* is both exciting and necessary, comparisons between studies are complicated by the use of different methodologies for assessing fat content, synthesis, and accumulation (*e.g.*, gravimetry, colorimetry, isotope studies, triacylglycerol kits; Tennessen et al., 2014). The use of distinct methodologies is problematic, because the range of available methods differ in reliability (Visser, LeLann et al., 2023; Williams et al., 2011) and in units of measurement, which can even differ within methods (absolute quantities, corrections with wet or dry weight, percentages etc…). When looking beyond our own studies on fat accumulation (using gravimetry; Enriquez et al., 2022; Visser et al., 2010, 2018, 2021) in the next paragraph, we are thus dependent on extrapolating general patterns based on comparisons within studies.

Our observation that *D. melanogaster* females show a net increase in fat content during adulthood is consistent with the literature. In our study, both lean and fat females increase their fat percentage between 7 and 14 days, but this increase was much more pronounced in lean females (+12.4%, with high values maintained until 28 days) than in fat females, which emerged with high teneral fat reserves (on average 22%), and showed a smaller increase (+4.2%) followed by a significant decrease at 21 days. Baenas & Wagner (2022) examined the effects of high-fat and high-sugar diets on triacylglycerol levels in *D. melanogaster* strain *w^1118^* over 10, 20, and 30 days. Female flies consistently maintained triacylglycerol levels ∼25% above controls on a high-fat diet across all three time points, and under a high-sugar diet from 20 days onwards, with no difference detected at 10 days on a high-sugar diet. Skorupa et al. (2008) further showed that dietary composition, rather than caloric intake, is a key determinant of triacylglycerol accumulation in adult *D. melanogaster*, with excess sugar strongly promoting fat storage and increased yeast availability decreasing fat storage. Indeed, flies maintained on the highest sugar-rich diet showed a progressive increase in triacylglycerol content up to 40 days of age, followed by a significant decline at very old ages (52-56 days). Taken together, these studies consistently showed that *D. melanogaster* females have a strong capacity to accumulate triacylglycerol throughout adulthood, driven primarily by sugar availability rather than by teneral fat reserves.

The late-life decline in triacylglycerol content observed by Skorupa et al. (2008) in flies that had accumulated large fat stores on a high-sugar diet, is consistent with the decrease in fat percentages we observe in our lean female flies at 49 days, after a peak at 14 days, suggesting that this late fat loss may reflect a more general feature in aging *D. melanogaster* females. The crossing of fat trajectories between lean and fat females, with fat females having a higher fat percentage during the first week of adult life but lean females surpassing them in fat content at 21 and 28 days, is consistent with studies showing that nutrient restriction during development can enhance fat storage capacity in adult *Drosophila*. Indeed, restricting food availability during larval development leads to higher fat reserves and greater starvation resistance in adult flies, an effect attributed to reduced insulin signaling that suppresses fat use during early adult life (Rehman and Varghese, 2021).

### **4.3.** Genotype and environment interact to determine patterns of fat depletion over time in L. heterotoma

In contrast to *D. melanogaster*, none of our five inbred lines of *L. heterotoma* showed an increase of fat content during adult life, regardless of whether wasps developed on lean or fat *D. melanogaster* hosts. This result differs from Visser et al., (2021), who reported fat accumulation in several outbred *L. heterotoma* populations reared on lean hosts. Several differences between the two studies should be considered. First, Visser et al., (2021) directly assessed host fat content and measured wasp free fatty acid synthesis using stable-isotope tracing, which allowed them to demonstrate *de novo* fatty acid synthesis and plasticity therein. In contrast, our gravimetric approach quantifies changes in whole-body fat content, meaning that the absence of fat accumulation in our inbred lines does not exclude that some fatty acid synthesis took place, but indicates that any synthesis was insufficient to compensate for fat depletion. Such a decoupling between fatty acid synthesis and net fat accumulation has been reported in many parasitoids (Ruther et al., 2021; Visser et al., 2021; Visser, LeLann et al., 2023). Combining our approach with stable-isotope tracing (as in Visser et al., 2021) would help disentangle net fat changes from underlying metabolic processes. Second, we used inbred lines derived from a single population, whereas Visser et al., (2018, 2021) used outbred field-caught populations: the genotypes represented by our inbred lines may lack the capacity for fat accumulation observed in other populations. Third, lean hosts were generated differently: Visser et al. (2021) used *D. simulans* that are naturally smaller and have a lower fat content, while lean *D. melanogaster* hosts were generated by manipulating the sugar content of the diet. Here, we used crowding to generate extremely small flies, so that our lean hosts contained even less fat (Enriquez et al., 2022; Visser et al., 2021). The absence of fat accumulation on these leaner hosts may also reflect the physical constraint discussed earlier (Visser et al., 2021; Visser, LeLann et al., 2023). Our results for wasps reared on fat hosts are, however, consistent with previous studies in which *L. heterotoma* reared on fatter hosts did not accumulate fat (2016 populations in Visser et al., 2018, 2021). Another, non-mutually exclusive explanation is that not only the quantity of fat is relevant, but also the quality of host-derived lipids. If the fatty acid composition of *D. melanogaster* is more favorable for use by the wasp compared to that of other host species, even a small amount of lipids carried over from the host could be enough to suppress adult wasp fat accumulation. Host identity, via lipid quality rather than quantity, may thus play an additional role in shaping adult fat dynamics in *L. heterotoma* and potentially other parasitoids (Thompson and Barlow, 1972, 1974).

Despite the absence of fat accumulation, we found a significant genotype x environment x age interaction for changes in fat content during adult female wasp life. Inbred lines differed in how quickly fat content declined with age, which is dependent on the phenotype of the host (lean or fat) and the inbred line in question. Previous work on *L. heterotoma* demonstrated plasticity of fatty acid synthesis in response to host conditions (Visser et al., 2021). Our results extend this finding by showing that the trajectory of fat depletion during adulthood varies among genotypes and developmental host conditions. To the best of our knowledge, such genetic variation in age-dependent fat-content trajectories has not been reported before in an adult parasitoid wasp (but see Reed et al., 2010 on genotype x environment interactions on fat accumulation in larvae). Genotype-specific rates of fat depletion can play an important role for life history trade-offs, because the rate of fat depletion will determine how long reserves last and, consequently, when energetic constraints on reproduction and survival set in (Arrese and Soulages, 2010). For now, it remains unclear whether such differences are adaptive. If such differences in the rate of fat depletion are maintained in natural populations, economizing on fat reserves may be less relevant during summer when host availability is high and longevity is expected to be lower in the field (due to *e.g.*, higher temperature, higher activity etc…). During winter, however, female *L. heterotoma* overwinter as adults in quiescence and we would expect that considerable lipid reserves are required for females to survive until spring (Enriquez and Visser, 2023). The rate of fat depletion may thus play an important role for overwintering, where a genotype depleting its fat reserves more slowly could have a higher chance of survival. To conclude, we showed that age-dependent changes in fat content during adulthood vary with genotype and developmental host conditions in *L. heterotoma*. The rate at which fat content declines with age thus depends on genotype, environment, and age, and could potentially evolve in response to selection.

## Acknowledgements

We are grateful to Patricia Gibert at the University of Lyon (France) for providing the *D. melanogaster* strain used in our experiments. We would also like to thank Dan Hahn for his feedback on an earlier version of this manuscript. This work was supported by the Fonds National de la Recherche Scientifique – FNRS under Grant(s) n° T.0186.20, J.0190.21, 40012205, and F.4570.25.

## Competing interests

No competing interests declared.

## Data statement

The data are available on Zenodo (https://doi.org/10.5281/zenodo.24305857).

## Notes

### Competing Interest Statement

The authors have declared no competing interest.

### Summary of Updates

This version of the manuscript has been revised following peer review. Title changed to better reflect what was measured. Terminology revised throughout: results are now described as age-dependent changes in fat content (fat accumulation and fat depletion) rather than fat metabolism. The Introduction has been updated to describe the host-parasitoid relationship and to compare our study more thoroughly with previous work on Leptopilina heterotoma. The Methods have been expanded to describe the experimental design, sampling and experimental unit in more detail. Statistical analyses have been revised: fat percentage is now analysed with models allowing group-specific variances. Discussion revised.

